# Millennial-scale societal shifts drive the widespread loss of a marine ecosystem

**DOI:** 10.1101/2024.05.19.594609

**Authors:** Sally C. Y. Lau, Marine Thomas, Jessica M. Williams, Ruth H. Thurstan, Boze Hancock, Bayden D. Russell

## Abstract

Degradation of marine ecosystems by human activities is a global problem, with only recent recognition that exploitation of ecosystems over millennia can result in their functional extinction and loss from human memory. To reconstruct the historical distribution of oyster reefs in China, and the context behind loss, we extracted information from archaeological records and historical documents (pre-modern Chinese literature, administration reports, art, maps, newspapers) spanning ∼7600 years, then constrained records with past coastlines and habitable environmental conditions. Oyster reefs were extensively distributed along >750 km of coastline in the Pearl River Delta, and their exploitation underpinned the region’s development into China’s first economic hub in the 6^th^ century. Millennial-scale overexploitation alongside societal shifts were central in their regional extirpation by the 19^th^ century, but the enduring cultural importance of oysters is maintained by aquaculture expansion. Informed conservation practices can be developed from reconstructing the temporal interplay between human societies and the natural environment.

## Introduction

Marine biodiversity hotspots are concentrated along the coasts of Southeast and East Asia (Jefferson et al., 2021) and are extremely vulnerable to, or have already experienced, widespread ecosystem degradation and destruction (Leadley et al., 2014; Wilkinson et al., 2006). In many cases, the loss of ecosystems has occurred through exploitation or expanding coastal development before their ecological and societal significance were understood (Wilkinson et al., 2006). Yet, understanding the ecological, cultural and economic context of a collapsed ecosystem is critical for successful conservation and restoration incentives (Gann et al., 2019). Unlike many parts of the world, the way these different contexts interact to contribute to the degradation of biodiversity hotspots across Asia is fundamentally unknown. As the destruction of coastal ecosystems continues, human memories of local resources can be forgotten within a few generations (Turvey et al., 2010). The undocumented loss of the physical ecosystems in the ecologically and culturally diverse Asian region could undermine the global understanding of both empirical ecology and the relationships between communities, land and sustainability (Caswell et al., 2020). Therefore, rebuilding the social-ecological history of the region can provide valuable insights that are required to reconcile and re-balance society’s relationship with nature, which could lead to more effective conservation and restoration actions (Gillson, 2022).

Globally, oyster reefs were historically a dominant structured ecosystem (Beck et al., 2011). The ecological role and the value of the associated ecosystem services that healthy oyster reefs provide are widely accepted (Grabowski et al., 2012; Smith et al., 2023; zu Ermgassen et al., 2020), as well as their historical significance in philosophical, ethical, and social aspects of many global coastal communities (Guillotreau et al., 2017; Wakefield & Braun, 2019). It has been estimated that at least 85% of global oyster reefs have been lost since the 19th century due to over-harvesting, habitat destruction and pollution (Beck et al., 2011). However, the global estimates of oyster reef loss and their societal contribution are likely underestimated as most data are derived from North American, European and Australian case studies. Very few natural oyster reefs have been documented across East and Southeast Asia (Beck et al., 2011; McAfee & Connell, 2020), but various lines of evidence indicate oyster reefs historically existed with strong cultural significance across the region. At least 16 native reef-building species can be found in the region (within the genera *Magallana* and *Tallonostrea*) (Sigwart et al. 2021), with the oyster harvesting and aquaculture industry dating back at least 800 years in countries throughout temperate and tropical Asia (e.g., Japan (Smith et al., 2018), China (Lam & Morton, 2003; Liu et al., 2021), Thailand (Chalermwat et al., 2003), Korea (Choi, 2008)). Archaeological records also highlight the widespread presence of middens with oyster shells (An & Lee, 2014; Engelhardt & Rogers, 1998; Hung & Zhang, 2019; Lee et al., 2017) and oyster-related cultural artefacts (Hung, 2019) throughout Asia, as well as the incorporation of oysters in architecture and ship construction materials in China and southeast Asia (Luengo, 2017, 2023; Yihang et al., 2021; Z.- J. Zhang et al., 2019). This anecdotal evidence regarding the historical prevalence of oysters introduces a major knowledge gap, as only a limited number of small oyster reefs are presently known to exist in China (Xu et al. 2023), with a complete absence of large-scale natural oyster reefs within living memory and modern literature.

Establishing evidence of the historical extent of oyster reefs in East and Southeast Asia is challenging as this region has experienced the world’s highest rate and extent of coastal modification. At least 92% of global coastal reclamation (i.e., filling the coastline to create additional land) has occurred within East and Southeast Asia, with most of the reclaimed area being within estuarine environments (Sengupta et al., 2023). Changes in land use patterns (e.g., clearing for agriculture, urbanisation), dam and reservoir construction, and changes to river flows are also common among many Asian estuaries and pre-date similar changes in western countries by centuries (Yunus et al., 2022). This results in sparse documentation of Asian oyster reefs in modern scientific literature, extremely limited official harvest records (cf. western countries), and scant records in the navigational charts. However, an alternative solution to reconstructing the historical significance of oyster reefs and their societal significance is to leverage the region’s cultural heritage. Many Asian countries, in particular China, are associated with documented human arts and history spanning thousands of years from ancient dynasties to colonialism and the modern era, offering insight into historical ecosystem baselines and degradation within the context of societal change.

Traces of human-ecosystem relationships, recorded in archaeological, written, and artistic forms, are increasingly being used to reconstruct the spatial and temporal extent of historical ecosystems where empirical records are depauperate (Erlandson & Rick, 2010; Günther, 1897; Mojetta et al., 2018; Thurstan et al., 2020). Here, we analysed archaeological, written and artistic evidence linked to oyster reefs in China over 7613 years, spanning the neolithic, bronze age, imperial China (1300 BCE - 1911 CE) overlapped by 16th - 19th century colonialism periods, through to the present day. We primarily focused on compiling historical documents relevant to the Pearl River Delta (PRD) in southern China (Fig. 1), as the pace and scale of change in the region, and consequent ecosystem alterations, provide a projection of large-scale biodiversity loss driven by globalisation. Additional evidence linked to historical oyster reefs in other parts of China are also presented. The Pearl River is China’s second largest river and its delta (∼9750 km^2^ in catchment area) is located in the sub-tropical zone in Southern China (Zhang et al., 2010), which borders Southeast Asia. The first local administration was established for the region during the Qin Dynasty (221-207 BCE) (Zong et al., 2013). Foreign trade began in the PRD ∼2000 years ago, with the region transforming as one of China’s leading port cities from ∼618 CE (Su & Grydehøj, 2022). Since 221 BCE, the PRD (and broader China) has experienced 12 imperial dynasty transformations (221 BCE - 1991 CE) (Wiebke et al., 2017), Portuguese (1557-1999 CE) (Amaro, 2016) and British (1841-1997 CE) (Henderson, 2001) colonialism and the modern era (People’s Republic of China). Today, ∼74 million people reside around the PRD (Fig. 1B), with the delta’s economy oriented towards the manufacturing industry.

**Fig. 1.**
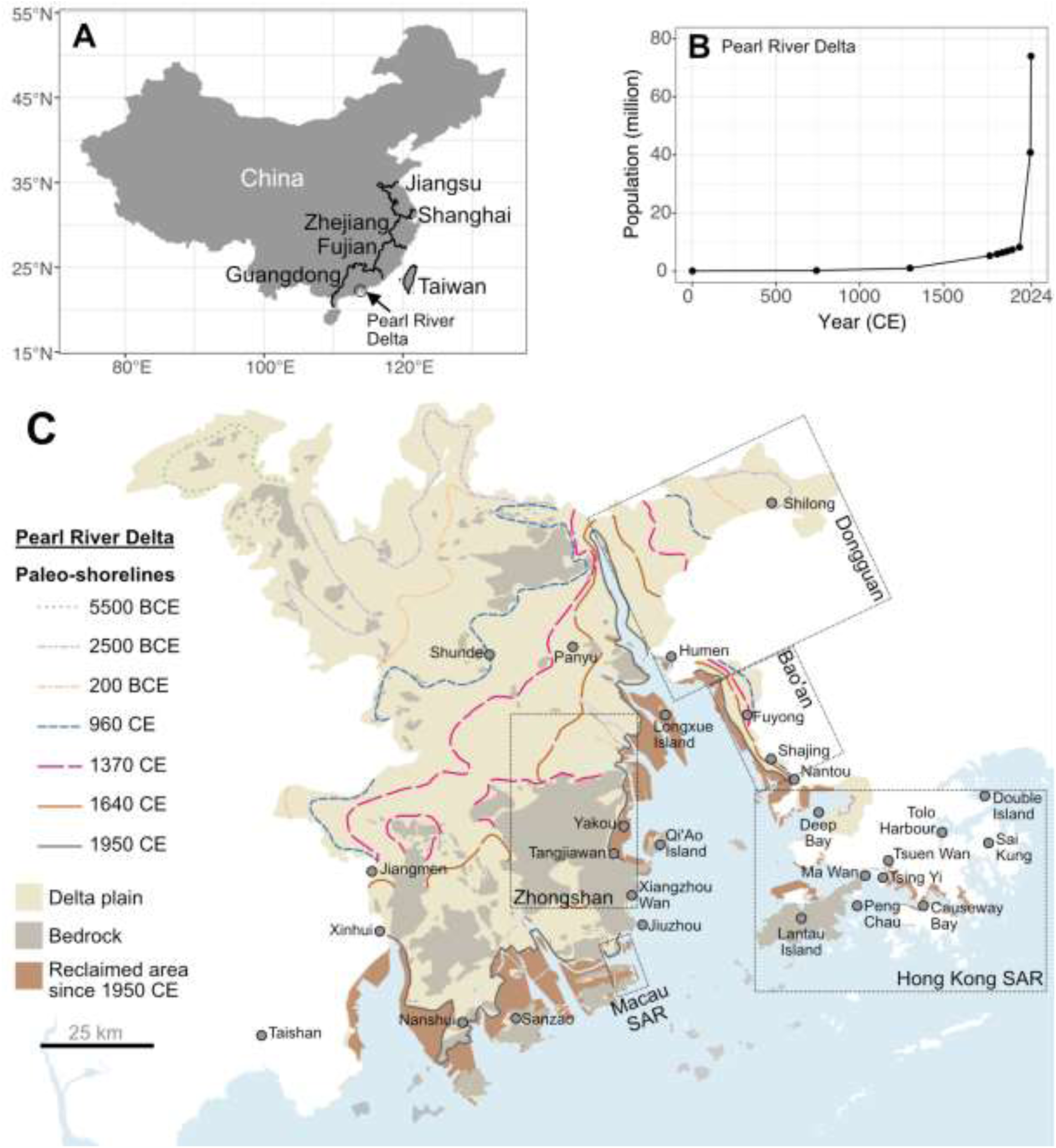
(A) Map of China with broad geographic localities where the presence of natural oysters has been mentioned in historical documents. (B) Human population size in the Pearl River Delta (PRD) between 2 and 2022 CE (EROS, 2024; Xiong et al., 2020). (C) Detailed map of PRD illustrating paleo-shoreline advances between 5500 BCE and 1950 CE, and locations (filled dots) where natural oysters have been mentioned in historical documents. Shoreline advances since 200 BCE until the present day are linked to ancient and modern human-induced land reclamation (Xiong et al., 2020). Reclaimed area = area of modern land reclamation since 1950 CE. Dotted box = geographically broader area relevant to the reported presence of oysters. SAR = Special Administrative Regions of China. Only modern town/county names are presented on map. Paleo-shoreline and map shapefiles were extracted from Xiong et al. (2020).

Evidence of ecosystem degradation also coincides with the development of the region ∼2200 years ago. The earliest changes include soil erosion from deforestation between 200 BCE - 960 CE leading to shoreline advances of 0.78 km^2^/year (Xiong et al., 2020; Zong et al., 2009). Between 960-1370, rapid shoreline advances of 2.69 km^2^/year coincided with a wave of human migration to PRD from central China due to wars between northern nomadic tribes and the Southern Song Empire (Xiong et al., 2020). Shoreline advances accelerated to 4.5 km^2^/year between 1370-1950, coinciding with a further increase in human population and the first documentation of land reclamation via physical trapping of sediments with dykes (Xiong et al., 2020). Since 1936, at least 84,300 hectares of additional intertidal and subtidal coastal marine area in the PRD have been reclaimed (Wei et al., 2021). Since the 1990s, the expanding population and economic boom (mostly from technology and manufacturing) led to sand mining for land reclamation, channel dredging and dam construction within the estuary (Wei et al., 2021). At least 89% of the natural PRD coastline are estimated to have been artificially modified through modern land reclamation (Zhang et al., 2022) since 1973. Recent human activities in the PRD have led to wholescale destruction of the remaining coastal habitats.

We hypothesise that natural oyster reefs historically existed in the PRD, meaning that degradation and loss of oyster reefs predates modern knowledge (e.g., Beck et al., 2011). This region is associated with the world’s oldest oyster aquaculture industry with oral history dating back to the 1300s (Lam & Morton, 2003; B. Morton & Wong, 1975). Today, the local perception of oysters in the PRD is mostly restricted to a declining aquaculture industry (Caswell et al., 2020), which is reliant on artificial substrates (Chan et al., 2022). The long history of human activities and ecosystem degradation alongside societal change and economic expansion is not unique to PRD; it is also common in other coastal areas in East and Southeast Asia that are habitable to oysters and other types of ecosystem engineers (Swerts & Denis, 2015). Therefore, documenting an ecosystem that is absent from modern knowledge, and understanding how the ecosystem was altered as a result of societal progression over time, would contextualise the breadth of loss of both biodiversity and the human-ecosystem connection within a global biodiversity hotspot.

## Results

### Past and present extent of oyster reefs in the PRD and broader China

There were a total of 101 document sources identifying the presence of shells and/or oyster habitats or reefs along the coastline of the South and East China Sea, including archeological records (n=4), modern journal articles (n=16), pre-modern Chinese literature (n=22 including three poetries), historical paintings (n=2), foreign travellers’ accounts (n=5), colonial government reports (n=4), archived newspaper articles written in modern Chinese (n=34) and in English (n=4), nautical map (n=1) and historical photographs (n=9). These records show that oysters were first reported along the coast of the PRD in 5590 BCE; and in broader China, in Guangdong since 806-820 CE, Zhejiang since 420-479 CE, Jaingsu and Fujian since 1020-1101 CE, and Shanghai since 1919 CE.

Historical documents indicated widespread oyster habitats were present in the PRD from the mid Holocene (∼5590 years BCE) until the early 1900s CE. The earliest shellfish records were observed through archaeological records in the form of shell middens and oyster shells in sediment cores, which were dated between ∼5590 BCE and ∼901 CE (Fig 2a). The distribution of the shell middens was extensive and found along the various PRD coastline configurations, following shoreline advancements (∼5500 BC - 960 CE) (Xiong et al., 2020). Based on historical records between 1600s and 1900s CE, oyster reefs were observed along 755 km of the main coastline of the PRD (Fig. 2b-f). As historical accounts also indicated the existence of subtidal oyster reefs (via documentation of extensive shell dredging), we infer the habitable area that historical oyster reefs (intertidal and subtidal) could have existed in the PRD to be 413,235 hectares (Fig 2g). In the modern era, however, subsequent scientific literature from the 1950s onwards indicates very few wild oysters could be found in the PRD.

**Fig 2.**
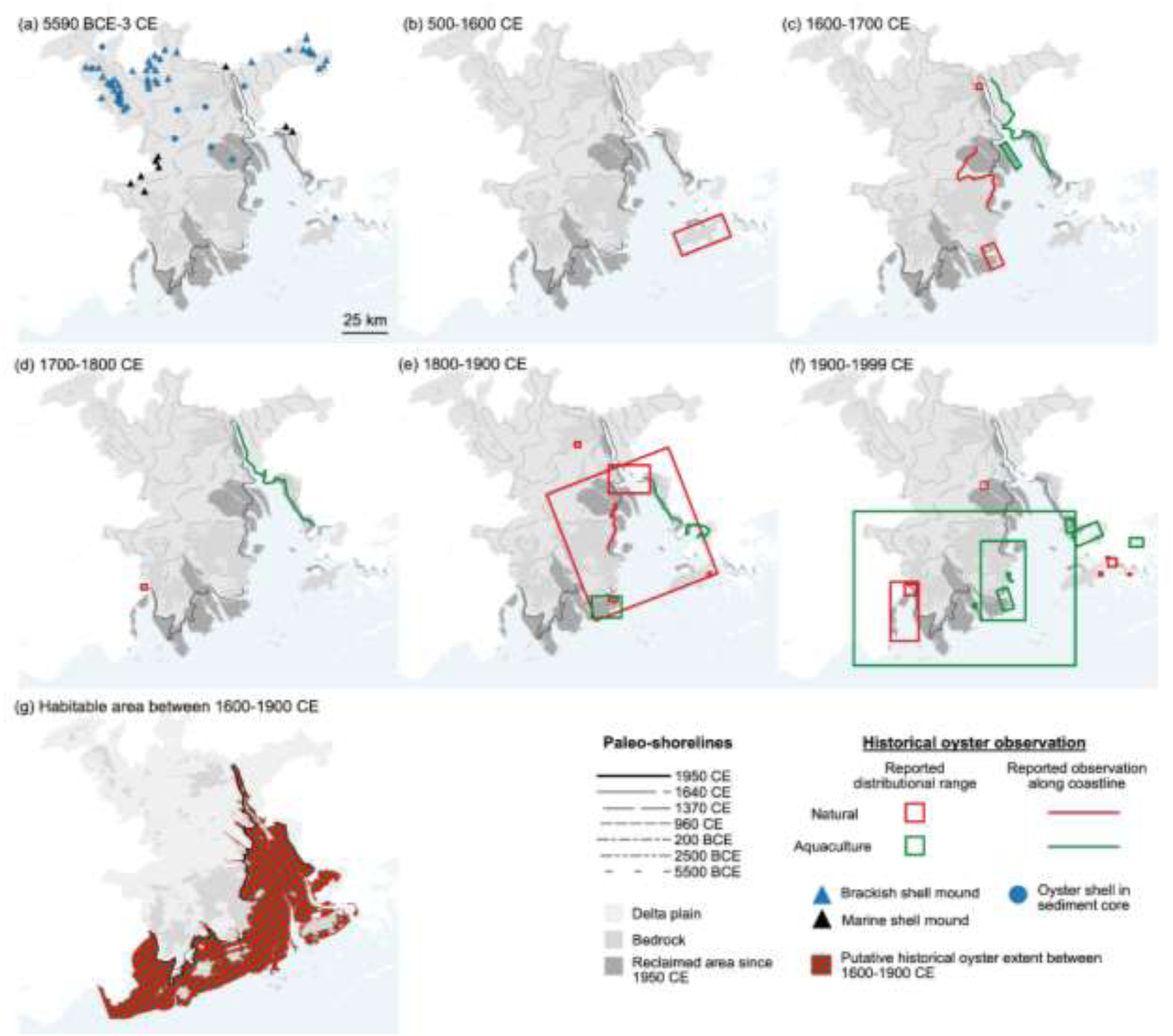
Reported distributional range of shell mounds and oyster shell in sediment cores from the Neolithic and Bronze Age (a), natural oyster observations and aquaculture observations between 500 and 1999 in the Pearl River Delta (PRD) (b-f), following paleo-shoreline distributions. (g) Habitable area that would have supported oyster reef formation between 1600-1900s in the PRD, following the 1950 (modern) PRD shoreline. Habitable area range limits were defined based on the intertidal and subtidal area between 0-12m of water depth, and between the furthest eastern to western points of past oyster observations, that is from Tsing Yi, Hong Kong, in eastern PRD to Sunning in western PRD.

### Overview of oyster reef descriptions in historical documents in the PRD

Early written records of oysters in the PRD are found in pre-modern literature written in literary Chinese, dated between 456 and 1741 CE. Of this pre-modern literature, seven localities were pinpointed to contain shellfish habitat, with seven additional sources describing the PRD region containing shellfish habitats (Table S1). Based on these descriptions, the PRD likely contained extensive oyster reefs; the descriptions of habitat extent and vertical relief included 1) “ expand in all four directions, with one or two zhang in height, precipitous like mountains” (Liu, 904) (1 zhang = ∼3 m), 2) “oysters grow on walls, up to 3 to 4 chi in height, can see them at low tide” (Qu, 1678) (1 chi = ∼10 inches), 3) “oyster, born in saltwater, grow on rocks, lumpy and connected together like houses, therefore each is called oyster house. Houses grow next to each other, spread outwards to dozens and hundreds of zhang.” (Li, 1803) (“hundreds of zhang” = minimum 300 m and up to kilometres).

Early foreign accounts on shellfish-human interaction in the PRD, including European explorers, diplomats, a missionary, a merchant and colonial government reports, were dated between ∼1400 and 1883 CE (Table S2). Of these early foreign accounts, Lantau Island (Schafer, 1957), Zhongshan (Bridgman, 1841) and Macau (Dabry de Thiersant, 1875) were pinpointed to contain shellfish habitats. An additional nine accounts also described the presence of shellfish habitats along the PRD coastlines. Between ∼1400 and 1922 CE, the PRD coastlines were also likely dominated by extensive oyster habitats. The accounts’ descriptions included 1) “Way to Canton: In the rivers and Channels there is taken abundance of Fish, Prawns, and the like, but particularly a vast quantity of Oysters” (Careri, 1700), 2) “In many parts of the river the poor find immense beds of oyster shells” (The London Saturday Journal, 1840), 3) “It is said that [shells] are thus procured, everywhere between bogue and Macao” (Martin, 1847), 4) “Notwithstanding the enormous consumption of oysters from time immemorial in China, there appears to be no diminution of the supply” (Dabry de Thiersant, 1875).

We also uncovered newspaper archives, written in English and traditional Chinese, on oyster-human interactions in the PRD, dating between 1881 and 1991 (see Fig. 4 for examples). From newspaper archives, a total of 19 locations were identified throughout the PRD, with nine locations reported with aquaculture activities and eight locations reported activities associated with the lime industry (Table S3). Early descriptions between 1881 and 1956 described depletion in natural oyster habitats, while later descriptions described productivity of local aquaculture industries (see section below), indicating a switch between wild harvest of declining natural oyster habitats to an increasing reliance on aquaculture production. Descriptions of natural oyster decline included: “Each spring villages harvest in the sea which erodes farmland nearby [in Panyu]” (Tsun Wan Yat Po, 1881), “An interesting local industry [Jiangmen] is the dredging of large shells” (Hong Kong Daily Press Office, 1909), “Riverways around Xijiang [in Xinhui]… oyster shells layer had always been on the river floor, not the oysters that are aquaculture annually… riverways should be around 1200 chi in length and 300 chi wide, but the actual areas covered in oyster shells, are now scarce” (The Kung Sheung Daily News, 1933) (1200 chi = ∼300 m), “Macau news: 43 oyster harvesters, seeking permission from Guangdong committee to harvest oysters in Zhuhai areas, to sustain livelihoods” (Ta Kung Pao, 1956).

Finally, prior to the type specimen of *M. hongkongensis* (species native to PRD) being formally described by Lam & Morton (2004), we found journal articles published between 1864 and 1983 (written in English), on shellfish-human interaction in the PRD. A total of 10 journal articles reported evidence of shellfish habitat in the PRD (Table S4). Four articles pinpointed that natural shellfish habitats existed around Hong Kong, including Peng Chau, Tsing Yi, Deep Bay and Causeway Bay (Schofield, 1983; Wong, 1984), which were associated with the lime industry in Hong Kong (Schofield, 1983; Wong, 1984). An additional two articles reported locations across the PRD being associated with the aquaculture industry, including Zhongshan, Macau, Kayo Island, Xin’an, Yuen Long and Deep Bay (Chan, 1935; Morton, 1976). From a synthesis of journal articles, the extent of these natural habitats by this time was likely limited as depletion was noted: “Practically all the public [aquaculture] oyster beds are in a depleted condition, scarcely producing sufficient oysters to make taking them worth while… There are plenty of natural beds, but they are all in an exhausted condition.” (Chan, 1935) “ lime is got entirely from coral and shell, and as the sea near it is almost worked out” (Schofield, 1983).

### Loss of natural oyster reefs and the emerging documentation of aquaculture

We found oyster reefs in the PRD sustained two types of industry, the lime industry (lime obtained from burning of shells) and aquaculture. Documentations of shell extraction for lime dated between 265-589 and 1933 CE (Table S5; n=20), and afterwards, the lime industry was only documented to exist on an isolated island (Peng Chau) until 1983 CE (n=2). Throughout the PRD, the lime industry likely operated from local to industrial scales, with documented methods ranging from burning of shells locally on the ground to use of large-scale kiln structures. Harvesting statistics are patchy and only found in sources from the late 1800s onwards. It should be noted that shell dredging had been documented in oral history dating back to 1403-1425 in the PRD (Hague, 1856) and paintings dating to 1637 in the broader China (Song, 1637), in relation to pearl oyster harvesting. Overall, harvesting efforts and scales were likely to be large across centuries. In 1741, a local government gazette documented that dozens of boats carrying oyster shells were overcrowding a creek in Xinhui (Qianlong Xinhui County Chronicle, 1741). One account also estimated hundreds of vessels were dredging shells in the PRD in 1847 (“everywhere between bogue and Macao”) (Martin, 1847). In 1883, 40 shell dredging vessels, with the smallest carrying ten tons, passed through a small creek in Shajing within half an hour (Hong Kong Daily Press, 1883). Similarly, in 1901, 95 shell dredging vessels (with Deep Bay as home port) were reported to operate within the vicinity of four-square miles (Hong Kong Government Gazette, 1901). The designs of shells dredging vessels around Shajing were of the same build, with dimensions around 40-50 feet in length by 10 feet in beam (Hong Kong Daily Press, 1883) (painting of a boat carrying oyster shells in Fig 3d).

**Fig 3.**
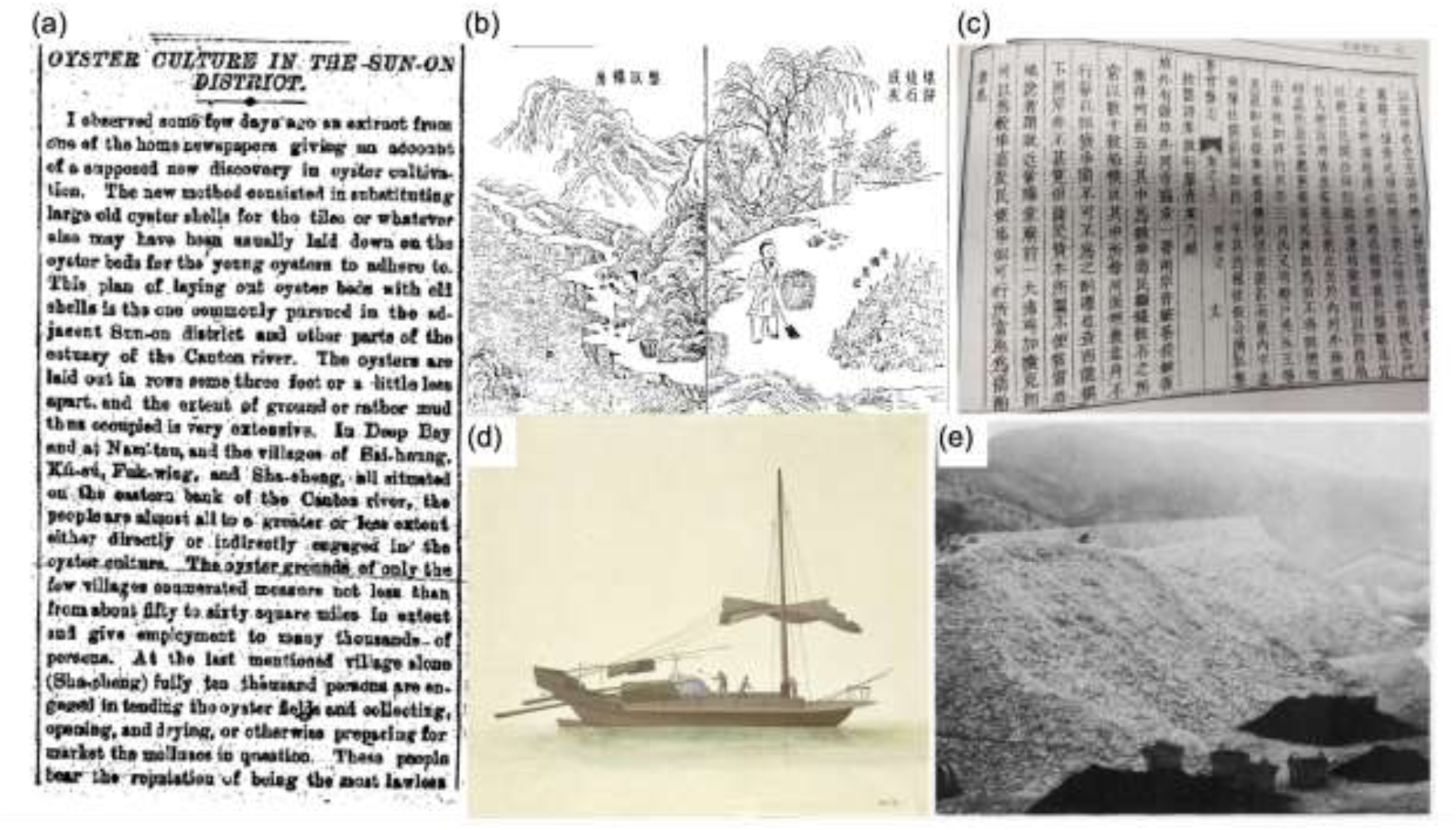
Representation of the variety of resources utilised to reconstruct historical oyster extent and cultural significance in the Pearl River Delta (PRD) and the broader China. (a) Excerpt of a newspaper article describing oyster aquaculture in PRD (Hong Kong Daily Press, 1883), (b) historical paintings of oyster shucking and shell burning on the ground for lime, depicted as the general usage of oysters in the broader China (Song, 1637), (c) an example of the documents used in the study – an excerpt of Chinese County Chronicle describing dozens of oyster shell boat blocking waterways around a loading port in Xinhui (Qianlong Xinhui County Chronicle, 1741)(for translation of excerpt used, contact authors), (d) painting of “Oyster Shells Boat 1800-1820” from the “Souvenir from Canton” collection (Qing dynasty Unknown Artist, 1800-1820) (tentative permission of use from the Victoria and Albert Museum; to be finalised prior to publication), (e) photograph of oyster shell heap collected from “seabed shellfish grounds” to be processed into lime in Tsing Yi, Hong Kong during 1950s (Wong, 1984) (permission of use from the Journal of the Royal Asiatic Society, Hong Kong Branch).

The earliest documentation of aquaculture in the PRD was dated to 1678, which was described as scattering rocks in the sea to form oyster beds around Dongguan and Xin’an (Eastern PRD) (Qu, 1678). These managed oyster beds were likely operating at an individual farmer level as the farmers were described as not allowed to cross each other’s beds (Qu, 1678), potentially indicating some early version of socially regulated farming leases. By 1883, the aquaculture industry was likely already at a large operational scale, with oyster grounds from Shajing reported to support ten thousand persons, with employment opportunities including collection, oyster opening and drying, and market preparation (Hong Kong Daily Press, 1883). By 1912-1921, it was reported that “nearly every bay and hamlet has [oyster] beds, from the Sunon district [East of PRD]… to Sunning [south of PRD]” (Order of the Inspector General of Customs, 1924). Between 1883 and 1991, oyster aquaculture had been reported throughout the PRD (Fig 2e-f). Given that these localities overlap substantially with earlier historical reports of the existence of natural oyster reefs, and that it spans the environmental conditions that promote oyster growth, it is likely that the increase in oyster aquaculture was driven by the ongoing demand for lime at a time when supply of oysters was declining from overharvest.

### Cultural imprints of oysters in the PRD and border regions

Finally, we found archaeological and written accounts describing the cultural influence of oysters across societies in the PRD. The earliest evidence of cultural influence of oysters in the PRD were described in archaeological excavations, where oyster shells were found as grave goods in prehistoric graves in the Bronze Age (Atha & Yip, 2016). Then, based on written documentations, oysters were first recorded as a form of currency in exchange for liquor in local markets, as well as used in local cuisines including stir fry and braising in the PRD in 888-904 CE (Liu, 904). A song was written in ∼1678 CE with lyrics describing oyster beds, as well as the actions of fisherwomen in harvesting oysters along the shorelines of southern PRD (Qu, 1678). In the PRD and its general province (Guangdong), oyster shells were also described as materials for wall building (sometimes referred as Chunambo) and window making in the 1600-1700s CE (Careri, 1700; Pinheiro et al., 2005; Qu, 1678; Zhaoqing government, 1726). In the broader China, lime from oyster shells was used for pest control in ∼1250 CE (Zhu, 1279), as well as during ∼1840-1875 CE for treatment of sores, fever, lack of appetites, skin inflammation, tumour and thrush (Dabry de Thiersant, 1875; The London Saturday Journal, 1840). Finally, we found illustrative paintings in ∼1637 CE depicting burning of oyster shells for lime and shellfish harvesting on shore in the broader China (Song, 1637), as well as of a junk carrying oyster shells in the PRD in ∼1800 CE (Qing dynasty Unknown Artist, 1820) (Fig 3d). We also found modern photographs taken in the 1870-1920 displaying oyster beds in Fujian and Shanghai (the broader China), and photographs taken in 1956-1964 depicting the remnants of industrial scale lime kilns (with oyster shells in the background) in Tisng Yi, Deep Bay and Peng Chau in the southern PRD (Fig 3e).

We discovered documentation indicating historical awareness of consequences from oyster loss in the PRD. Two petitions dated back to 1931 in Shunde in the upper PRD and 1933 CE in Xinhui in the western PRD, which requested local authorities to cease shell harvesting. The reason stated in the 1931 petition was “to improve livelihoods” (The Chinese Mail, 1931). The petition from 1933 indicated that locals (in Xinhui, PRD) deduced natural oyster reefs could prevent local flooding; and a campaign was launched to cease shell dredging to prevent damage from flooding in local villages (The Kung Sheung Daily News, 1933). A poem dated back to the ∼1100 CE outlined the biodiversity associated with oyster reefs in Fujian, describing ∼23 species living within oyster crevices including snails, crabs, clams, jellyfish, cuttlefish, softshell turtle and fishes, and that “people with long handle nets” were putting these “delicious produce” in danger (Liu, 1147). Overall, these sources demonstrate a community awareness that oyster reefs historically formed a natural part of the PRD (and broader China’s) ecosystems, the ecosystem services that these reefs provided (from shoreline protection to fisheries provisioning) and, importantly, the community-wide concern of their decline and loss. A timeline of all events on historical oyster presence, lime industry, shell dredging, aquaculture and their cultural influence are displayed in Fig 4.

**Fig 4.**
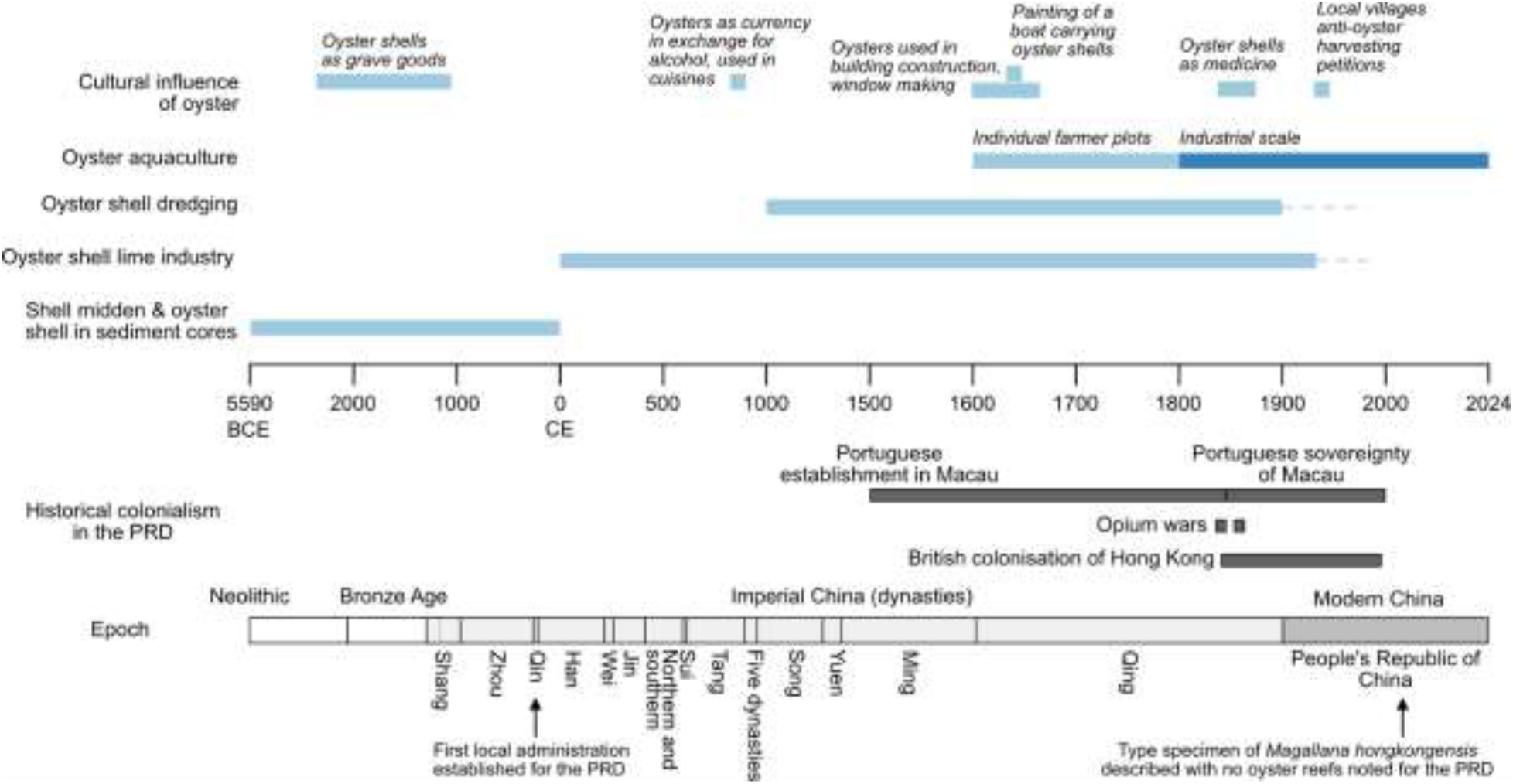
Timeline of events extracted from historical records on the historical presence of oysters in Pearl River Delta (PRD), the dating of oyster-related industries, and the dating of cultural influence of oysters. Dash lines representing only one location, an isolated island in Hong Kong (Peng Chau), was mentioned for oyster shell lime industry and dredging during the time period. Timeline of historical colonialism in the PRD was extracted from Amaro (2016) and Henderson (2001). Timeline of Imperial China dynasties was extracted from (Wiebke et al., 2017).

## Discussion

The historical documentation that we uncovered indicates widespread oyster reef ecosystems with broad economic and cultural significance once existed in the Pearl River Delta (PRD) in Asia until the late ∼1800s. The PRD has experienced continuous shoreline advances from land reclamation since 200 CE, with almost the entire PRD shoreline, tidal current regime and sediment load being modified throughout the centuries until the present day (Xiong et al., 2020). Since 1973, 89% of the natural PRD coastline has been modified through modern land reclamation (Zhang et al., 2022), and the habitats where native oyster reefs would have existed and benefited the PRD’s society for the last millennium no longer exist. Historical observations pinpointed natural oyster reefs throughout the PRD until the early 1900s. Between the 1400s and 1980s, oysters were harvested via dredging and burnt as raw material for the once prosperous lime industry. Oyster harvesting in the PRD was extensive both in space and time and ecological evidence of oyster reefs was physically removed. The historical documentation tells the tale of abundant reefs exhibiting a strong cultural influence (grave goods, cuisines, currency, medicinal treatments, poetry, songs, paintings, building constructions and village petitions) since the bronze age until 1930s. In contrast, the modern literature (1950s onwards) describes the absence of natural oyster reefs in the region (Lam & Morton, 2004). The environment once inhabited by oyster reefs now largely exist in the form of bare soft-sediment substrate, densely stocked aquaculture leases, or as extremely degraded, sparse, and spatially limited natural oyster beds in very few intertidal areas in the PRD (e.g., in parts of Hong Kong; Lau et al., 2020). This study, therefore, establishes the first baseline demonstrating that extensive natural oyster reef with vertical relief did historically exist in Asia. The spatial and temporal coverage of the documentation we have uncovered further makes it possible to identify the historical extent of these reefs and their previously underestimated importance in society.

### Cultural context of oyster loss through the millenia

The existence of oyster reefs in the PRD (and across Asia in general) has been overlooked and under-recognised in modern scientific and history literature; constituting a major data gap (i.e., the almost complete lack of data from Asia in Beck et al. (2011)). Our findings, however, show that the now collapsed oyster reefs once sustained a large-scale lime industry, which would have provided essential raw materials to position the PRD as China’s economic hub from ∼618 CE. For example, sticky rice-lime mortar (derived from shell lime) was a common construction material in Northern and Southern dynasty (420–589 CE) (Yang et al., 2009) through Tang, Yuen, Ming and Qing (1644–1911 CE) dynasties (Shao et al., 2019; Weng et al., 2021; Wu, 1989; Yang et al., 2009). The first documentation of oyster shell dredging (∼1403 CE) also roughly followed a human migration wave in the PRD in the 13th century; by the end of the 13th century, ∼85% China’s inhabitants were believed to live in southern China due to Mongol conquest during Yuan dynasty (Bai, 2022). The economic legacy derived from oyster utilisation and the early population expansion remains to this day but seems to have been eclipsed by the perception that the region’s first economic boom was the industrial and technology boom that began in ∼1970s (Su & Grydehøj, 2022). Oysters also historically had a strong cultural influence on the PRD’s society, ranging from grave goods to architecture designs, as a form of currency, the topics of songs, poetries, paintings, the representation of women, and the basis for cuisines and medicinal purposes. Interestingly, the importance of oyster reefs in the region is also exemplified as the subject of conservation campaigns first dated from the bronze age but continuing through to the present day.

In terms of society and social relations, collating literature on oyster-human cultural interactions around the PRD, reveals that people whose livelihoods depended on oyster extraction and/or cultivation (including fishers and lime workers) were likely historically to be perceived as “social outcasts”, strongly contrasting the perceived importance of oysters. In other parts of the world, groups who harvested fisheries resources, such as oysters, were not marginalised from societies (Humphreys et al., 2014; Schulte, 2017). In southern China and the PRD, elements of marginalisation were found in folklore dating back to 1178 CE, where people who inhabited offshore islands ate oysters and built houses with shells were referred as “Luting”, descendants of Jin dynasty outlaws (265-420 CE), and described as barbaric naked beasts with cone-shaped heads, and to “play in the waves like otters” (Fung, 1998). Over the centuries, fishers in the PRD were generally associated with having a lower social and legal status (Fung, 1998; Watson, 2022), right up until the 1950s, where the concept of piracy was still associated with oyster harvesting in Hong Kong (B. Morton, 1975). One account in 1883 described the PRD oyster cultivators bared “ the reputation of being the most lawless and turbulent in the province”, labelled as “broken lives” locally, and were “holding human life at a very cheap rate indeed, either their own lives or those of other people” (Hong Kong Daily Press, 1883). It is thus probable that oyster-associated activities could be underrepresented in historical records and accounts due to social biases (McCullagh, 2000). It should also be noted that few records are likely to have survived throughout dynasty regimes, wars and changes in governments; for example, the burning of books and loss of records to fire have been documented during imperial China across dynasties (Brokaw & Chow, 2005; Petersen, 1995; Wang & Zhang, 1988). Yet, we show here the value of historical documents in refining the global underestimation of biodiversity and human-ecosystem connection loss – perhaps even more so in regions where documentation is sparse, and no knowledge of ecological baselines exists. Through understanding the historical ecology of oyster reefs in the PRD, we define the livelihoods of the local communities that once sustained the economic growth of the PRD over the millennia.

At least 85% of global native oyster reefs have been lost largely due to over-harvesting during the 19th century (McAfee & Connell, 2020), which is also one of the main causes of oyster reef collapse in the PRD alongside the known extensive shoreline modification (Xiong et al., 2020). Recent advances in historical ecology have recognised the sustainable historical indigenous practices of harvesting and managing oyster reefs prior to colonialism, and have advocated for the inclusion of indigenous knowledge in modern oyster reef management (Reeder-Myers et al., 2022). In contrast, the case of the PRD does not necessarily follow this narrative. We documented a continuous dynamic relationship between oyster reefs and human livelihoods, which were shaped alongside the challenges of dynasty transformations, colonialism, and cultural, societal and economic progression throughout the last two thousand years (Fig. 4). This begins to highlight the fundamental challenge of balancing cultural heritage and ecology with economic progression and industrial development. We anticipate this is not a case unique to the PRD, but rather applicable to other places in Asia, as evidenced by preliminary syntheses of historical case studies from Japan, Korea and China (Williams et al., 2024).

### End of shell harvesting in the wild followed by the expansion of aquaculture

As natural oyster reefs in the PRD began to disappear from historical records in the late 19th century, methods of harvesting transitioned from wild harvest fisheries to oyster aquaculture. We found that oyster aquaculture began to be mentioned at higher frequency through time; beginning from small-scale individual aquaculture farms in 1678, to industrial scale (i.e., farms supporting >1000 jobs per town) being reported in 1883. Today, the oyster aquaculture industry, although declining, still exists around Deep Bay in the PRD (Caswell et al., 2020). Outside of aquaculture zones, natural recruitment and degraded and patchy wild oyster beds consisting of native species have been observed around Lantau Island and Tolo Harbour of Hong Kong (Lau et al., 2020). These areas lack the hard substrate required for the active growth of reefs (Hemraj et al., 2022), combined with ongoing coastal development and pollution (Hong et al., 2021), natural oyster reefs have not re-established since industrial-scale harvesting stopped in the 19^th^ century. Without these reefs, we hypothesise that the source of oyster larvae is likely to be the relatively extensive traditional benthic oyster farms, which have been abandoned for decades (Chan et al., 2022). In contrast to modern oyster raft techniques that often selectively culture triploid oysters in the PRD aquaculture zones, these abandoned farms relied on natural recruitment to farm structures. Left unmanaged for decades, these structures now support mature oyster clumps that can act as not only brood stock, but also some of the functions and services of natural reefs (Chan et al., 2022).

### Modern conservation of Asian oysters through reconciling historical ecology and lost social history

Historical documents collectively revealed ∼755 km of coastline, encompassing 413,235 hectares of habitable area, in the PRD where historical oyster reefs could have existed between 1600-1900s CE. Oyster reefs were likely to have been common across this region and a key component of the broader habitat mosaic, complementing and protecting seagrasses, mangroves and soft sediment shores natural within the PRD (Sun et al., 2021). Despite this historically broad geographic distribution, restoration on this scale under contemporary environmental conditions is not a realistic target, as the sediment regime (Zhang et al., 2012), channel morphology (Zhang et al., 2015) and substrate availability (Lau et al., 2020) in the modern PRD has been altered through ongoing human activities.

To re-integrate oyster reefs into modern ecosystem restoration efforts in the PRD, practitioners should first focus on restoring realistic and probably smaller scale oyster habitats. Prior to this current study, however, it was not even recognised that oyster reefs were both an extensive ecosystem across the region, and also intricately woven into the culture of local groups, whose culture endures to the present day (e.g., villages and families that have practised aquaculture for generations). By longitudinally documenting the social-cultural history around the reefs, we provide a historical foundation for current and future generations to reconcile their relationship with nature and offer lessons regarding the future management of such ecosystems for practitioners and communities. By extension, we also set an example for other studies seeking to engage modern societies with these important components of the nearshore ecosystem. Restoring the lost local history and culture can offer a new dimension for ecosystem restoration.

## Methods

Contemporary field campaigns in the lower Pearl River Delta (PRD) region (Lau et al., 2020), published sediment cores (Zong et al., 2009), and trawling and grab sampling records (Shen et al., 2022; Z. Wang et al., 2021; Wu & Richards, 1981; Zeng et al., 2012) obtained throughout the PRD, as well as the known extent of coastline reclamation (Wei et al., 2021; X. Zhang et al., 2022), indicate that there is unlikely any remaining physical evidence of natural oyster reefs in the region. Therefore, we systematically reviewed the scientific literature and historical documents to evaluate the historical existence of oyster reefs. Our primary geographical focus was the PRD region, but additional evidence linked to historical oyster reefs in the broader Guangdong and other parts of China was also collated when possible. Throughout March 2018 and July 2023, a keyword search of combinations varying between “Zhujiang”, “Pearl River Delta”, “Pearl River Estuary”, “Pearl River”, “Kwantung”, “Canton”, “Hong Kong”, “Macau”, “Macao”, “Zhong Shan”, “Guangdong”, “oyster”, “reef”, “lime”, “shell”, “shellfish” in English, and simplified and traditional Chinese was conducted on online databases and electronic document repositories. Note that in the English language, Pearl River Estuary, Zhujiang, Pearl River, Canton are synonyms and/or historical names for the Pearl River Delta; Kwantung is synonym and/or historical name for Guangdong; and Macau and Heukang are a synonym and/or historical name for Macao. Databases and electronic document repositories included the Google Search Engine (https://www.google.com/), Google Books (https://books.google.com/), Google Scholar (https://scholar.google.com/), HathiTrust Digital Library (https://www.hathitrust.org/), Internet Archive (https://archive.org/), JSTOR (https://www.jstor.org/), Biodiversity Heritage Library (https://www.biodiversitylibrary.org/), the Chinese Text Project (https://ctext.org/), Taiwan National Central Library Special Collections Digital Images (https://rbook.ncl.edu.tw/NCLSearch/home/info), Multimedia Information Service of the Hong Kong Library (https://mmis.hkpl.gov.hk/) and the University of Hong Kong library archive (https://lib.hku.hk/). All curated resources are available on request to the authors.

Extracts of documents that mentioned oysters were recorded, and relevant locations mentioned within the text were pinpointed at the highest geographic resolution possible on the modern map of the PRD (as of July 2023), ranging from towns or defined positions on the coast (highest resolution) to the broader PRD region (lowest resolution). Historical town names and locations mentioned were matched with historical maps from the same time period to pinpoint the correct coordinates for the locations. For documents without locations explicitly stated (e.g., poetry), we used these resources to indicate 1) the cultural significance of oysters (i.e. non-provisional uses, symbolism), and 2) the general size and location of oyster reefs in broader China.

### Estimation of historical oyster reef extent

To estimate the maximum potential area and length of coastline that historically had oyster reefs as an ecological habitat, we traced the potential historical oyster reef extent based on locations and observed range mentioned in the collated documents following the paleo- and modern-shoreline configurations of the PRD in Xiong et al. (2020). For specific areas (e.g., Peng Chau Island in Hong Kong), we estimated the oyster reef extent along the shoreline of Peng Chau Island. For a specific larger area such as county (e.g., Zhongshan County), we estimated the oyster reef extent along the shoreline of the county boundary of Zhongshan. For a specific range with two endpoints (e.g., “between Bogue and Macao”), we estimated the oyster reef extent along the shoreline between the two locations. For oyster extent estimation within a specific time period (Fig. 2), we used the shoreline configuration from the available time period that is closest to the date mentioned in the source. If an area or range was mentioned by multiple historical accounts, the area was only measured once.

We also quantified the area that would have been habitable by oyster reefs between 1600s and 1900s using QGIS version 3.22 (QGIS.org, 2024), following the 1950 CE configuration of the PRD shoreline. We first defined the range limits of historical oyster reef extent across the PRD based on the furthest eastern to western points of past oyster observations, that is from Tsing Yi in Hong Kong to Taishan (Sunning) in western PRD. Since the maximum known subtidal water depth limit of the locally found *Magallana* species (*M. gigas, M. ariakensis* and *M. hongkongensis*) in PRD range to 10-13 m (Meng et al. 2018; Quan et al. 2013; Lam & Morton 2004), we infer the habitable areas for historical oyster reef would have existed from the intertidal (0 m) to subtidal area up to 12 m between Tsing Yi in Hong Kong to Sunning in western PRD. This 12 m lower depth boundary also coincides with the salinity regime up to ∼30 psu in the PRD (Zhi et al., 2022), which is a threshold that is tolerable for *M. gigas*, *M. ariakensis* and *M. hongkongensis* (Huo et al., 2014; Qin et al., 2020; Zhao et al., 2012). High resolution, point-based water depth information of the PRD was extracted from Zhang et al. (2021). The point-based water depth data were then interpolated using Triangulated Irregular Network (TIN) interpolation in QGIS, and area extent was extracted by clipping the seafloor area between Tsing Yi to Sunning from 0 m (based on PRD 1950 CE shoreline configuration) to 12 m water depth. It is worth noting that this estimate is likely to be a conservative estimate as other subtidal reef-building species possibly also inhabited the area (Lam & Morton, 2004), but for which there is only limited historical evidence of reefs at greater water depths in the PRD (e.g., *Ostrea* spp.). In addition, the eastern waters of Hong Kong (e.g., Tolo Harbour) are more influenced by the South China Sea (higher salinity) than the PRD so have a different oyster species assemblage; even though we found historical records of oyster reefs in the eastern region we did not consider the eastern waters of Hong Kong in the calculation of potential habitable area as records were more sparse and less accurate in locations than for the PRD.

## Acknowledgements

We thank Jun Cheng for discussion about oysters in China, 劉偉𤋮 and 林詠芝 for their invaluable help with translation, Alyssa Chan for document organisation, and Khan Cheung, Jacky Chun, Timothy Yin, and Donald Chan for help with documents.

